# Graph Peak Caller: calling ChIP-Seq Peaks on Graph-based Reference Genomes

**DOI:** 10.1101/286823

**Authors:** Ivar Grytten, Knut D. Rand, Alexander J. Nederbragt, Geir O. Storvik, Ingrid K. Glad, Geir K. Sandve

**Author notes:** These authors contributed equally to this work (Ivar Grytten, Knut D. Rand).

## Abstract

Graph-based representations are considered to be the future for reference genomes, as they allow integrated representation of the steadily increasing data on individual variation. Currently available tools allow *denovo* assembly of graph-based reference genomes, alignment of new read sets to the graph representation as well as certain analyses like variant calling and haplotyping. We here present a first method for calling ChIP-Seq peaks on read data aligned to a graph-based reference genome. The method is a graph generalization of the peak caller MACS2, and is implemented in an open source tool, *Graph Peak Caller*. By using the existing tool *vg* to build a pan-genome of *Arabidopsis thaliana*, we validate our approach by showing that Graph Peak Caller with a pan-genome reference graph can trace variants within peaks that are not part of the linear reference genome, and find peaks that in general are more motif-enriched than those found by MACS2.

**Author summary:** The expression of genes is a tightly regulated process. A key regulatory mechanism is the modulation of transcription by a class of proteins called transcription factors that bind to DNA in the spatial proximity of regulated genes. Determining the binding locations of transcription factors for specific cell types and settings is thus a key step in understanding the dynamics of normal cells as well as disease states.

Binding sites for a given transcription factor are typically obtained through an experimental technique called CHiP-seq, in which DNA binding locations are obtained by sequencing DNA fragments attached to the transcription factor and aligning these sequences to a reference genome. A computational technique known as *peak calling* is then used to separate signal from noise and predict where the protein binds.

Current peak callers are based on linear reference genomes that do not contain known genetic variants from the population. They thus potentially miss cases where proteins bind to such alternative genome sequences. Recently, a new type of reference genomes based on graph representations have become popular, as they are able to also incorporate alternative genome sequences. We here present *Graph Peak Caller*, the first peak caller that is able to exploit such graph representations for the detection of transcription factor binding locations. Using a graph-based reference genome for *Arabidopsis thaliana*, we show that our peak caller can lead to better detection of transcription factor binding locations as compared to a similar existing peak caller that uses a linear reference genome representation.

## Introduction

Transcription factors are known to play a key role in gene regulation, and detecting regions associated with transcription factor binding is an important step in understanding their function. The most common technique used to detect transcription factor binding sites is *ChIP-seq*, combining chromatin immunoprecipitation (ChIP) assays with sequencing. Obtaining putative binding regions from ChIP-seq data is done using computational techniques known collectively as performing *peak calling*. Several *peak callers*, programs to perform peak calling, have been developed for this purpose, for example MACS2 [1] and SPP [2] [3]. Common for all current peak callers is that they take reads mapped to a *linear* reference genome, such as GRCh38, as input.

Graph-based reference genomes offer a way to include known variants within a population in the reference structure [4]. The software package *vg* supports mapping reads to a graph-based reference genome with potentially increased accuracy [5, 6] as compared to mapping reads to a standard linear reference genome using tools like BWA [7] or Bowtie [8]. Several types of genomic analyses, such as variant calling and haplotyping, can now be performed using graph-based references [5, 6]. However, no tool currently exists for performing peak calling on graph-based references.

## Results

We present *Graph Peak Caller*, a first method for detecting transcription factor binding events from ChIP-seq reads mapped to a graph-based reference genome. Graph Peak Caller is based on the same principles used by MACS2 (see Fig 1 for an overview), and is able to call peaks with or without a set of control alignments. For the case of a graph that merely reflects a linear reference genome, our peak-caller produces the same results as MACS2. As input, it supports alignments in the Graph Alignment/Map format (GAM) from *vg*, as well as reads represented as genomic intervals using the Offset Based Graph Python package [9]. Graph Peak Caller can be run from the command line, and is also available through Galaxy at https://hyperbrowser.uio.no/graph-peak-caller. In the Github repository at http://github.com/uio-bmi/graph_peak_caller, we provide a simple tutorial on how to use *vg* and Graph Peak Caller to go from raw ChIP-seq reads to peaks.

**Fig 1.**
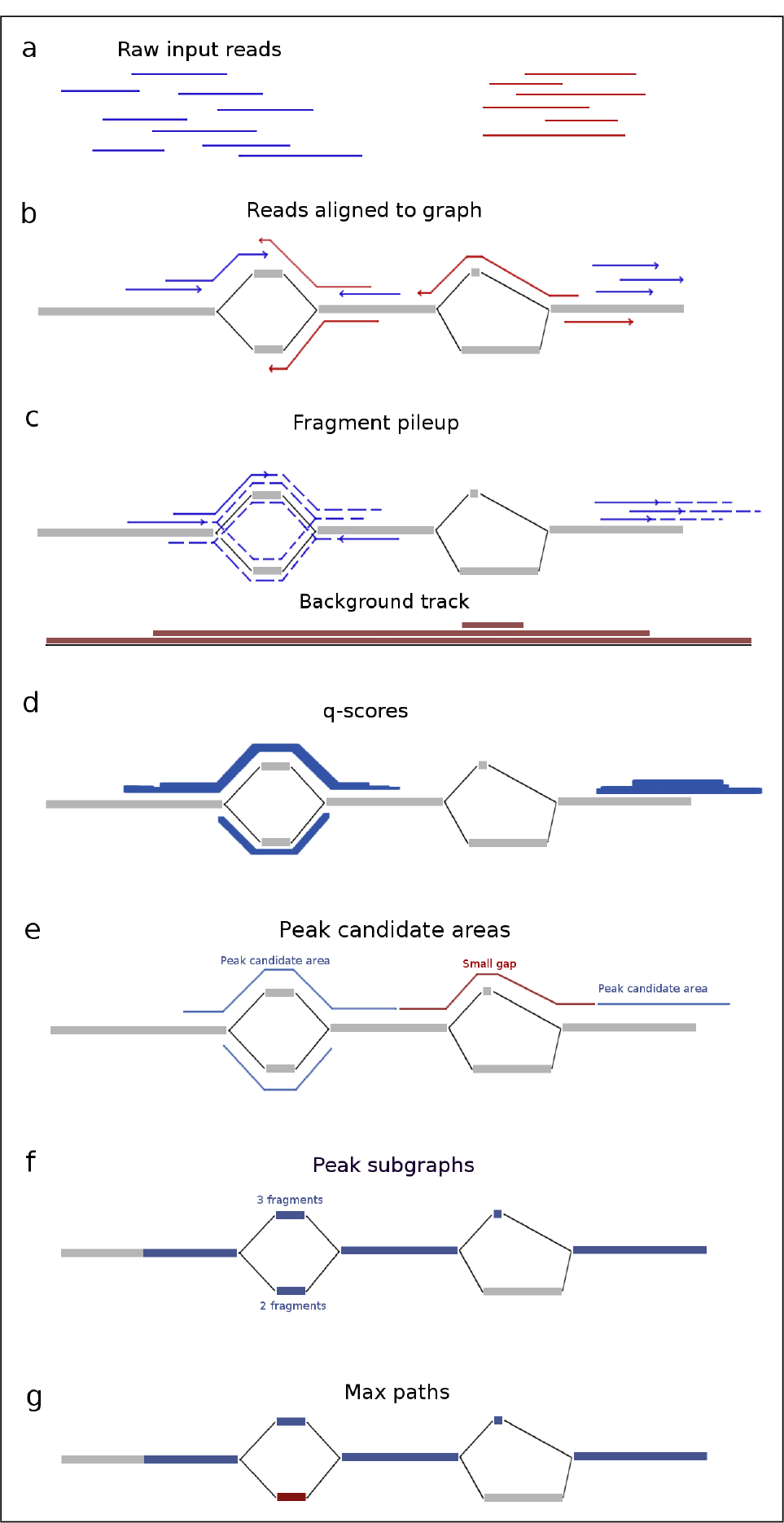
Overview of how Graph Peak Caller works. Example of peak calling on an example graph (nodes in gray, edges in black). After raw reads (a) in the form of input reads (blue) and control reads (red) have been mapped to the graph-based reference genome and filtered based on mapping quality (b), the fragment pileup is created (c) by extending the forward input alignments and reverse input alignments (extensions shown as dotted lines) along all possible paths in their corresponding direction. A background track is created by projecting the alignments resulting from the control reads onto a linear path and calculating a local average of read counts. Then the linear track is projected back to the graph again (not shown). The fragment pileup is treated as *counts* and the background track as *rates* in a Poisson-distribution, and p-values are computed for each position for the observed count, given the corresponding rate. Adjusted q-values are computed to control the false discovery rate (d) (figure shows q-scores, which are −*log*_10_(*qvalue*)). The q-values are thresholded on a user-defined threshold (default 0.05), resulting in a set of peak candidate areas with gaps between them (e). Small gaps are filled, resulting in a set of peak subgraphs (connected subgraphs)(f). Graph Peak Caller finds a single “maximum path” (g, maximum path in blue) through each peak subgraph by selecting the path that has the highest number of input reads mapped to it.

The output from Graph Peak Caller consists of graph intervals, but the tool is also able to transform these into approximate positions on a linear reference genome (by projecting them to the nearest position on the linear reference genome), making it possible to analyse detected peaks further using existing “linear” approaches. Graph Peak Caller is also able to output peaks candidates for differentially expressed peaks.

To showcase and test Graph Peak Caller, we chose to perform peak calling on *Arabidopsis thaliana*, as the 1001 Genomes Project for *A. thaliana* makes it possible to build a high-quality reference graph with a high density of variants (on average one SNP or indel for every 9 base pairs, compared to one SNP or indel for every 27 base pairs in the human 1000 Genomes Project).

We called peaks on a graph-based reference genome for *A. thaliana* and compared the results to peaks called on the *Tair10* [15] linear reference genome by MACS2 (Methods).

Table 1 shows an overview of peaks found by Graph Peak Caller and MACS2. Most of the peaks found by one peak caller are also found by the other. Among these, the peaks found by Graph Peak Caller are slightly more enriched for DNA-binding motifs than the peaks found by MACS2 for all transcription factors, except SEP3, where the numbers are the same. Fig 2 shows an example of a peak detected on chromosome 1, illustrating how both peak callers find a peak, but only the peak found by Graph Peak Caller has a match against the motif.

**Table 1.**
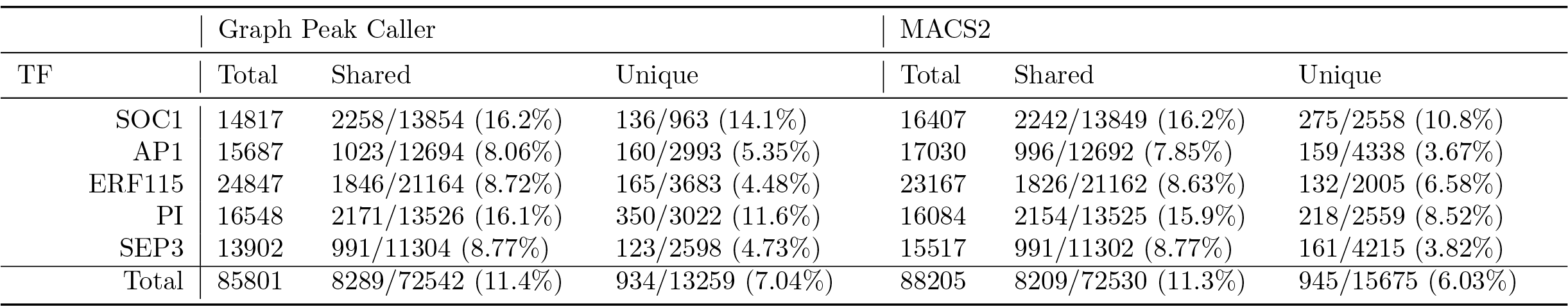
Overview of peaks reported by Graph Peak Caller and MACS2 on *A. thaliana* for 5 transcription factors (TFs). *Total* is the total number of peaks reported by the peak caller, *shared* is the number of peaks that overlap with one or more peaks from the other peak caller, and *unique* is the number of peaks reported by one peak caller and not the other. In the categories *shared* and *unique*, both the number of peaks with motif match (the number before the /) and the number of peaks found are shown (percent of peaks with motif match are shown in parentheses). Here, all peaks have been trimmed to 120 base pairs around the peak summit (position in peak with lowest q-value), to make the comparison clearer.

**Fig 2.**
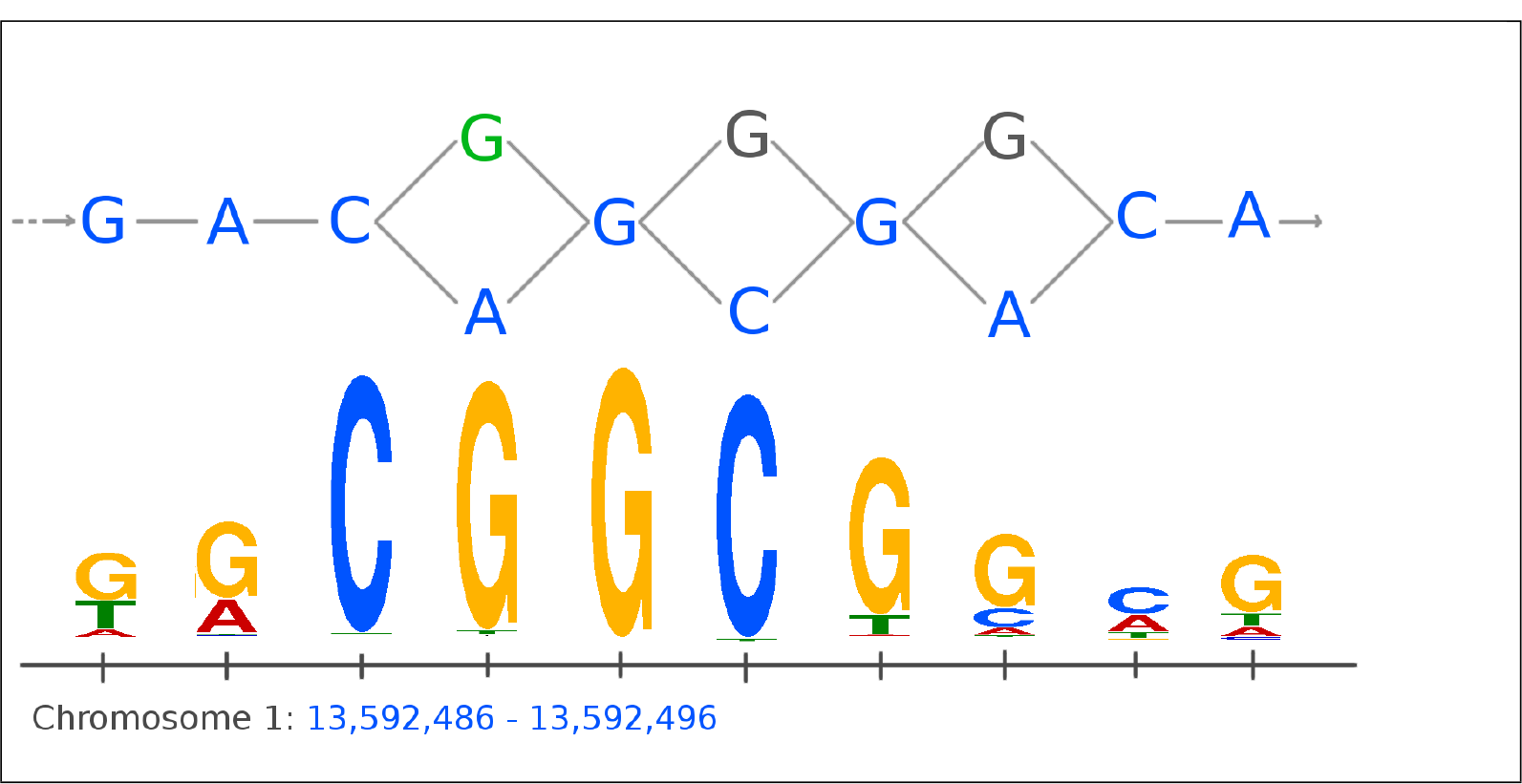
Example of a motif match for a peak following a variant not part of the linear reference genome. Showing part of the graph-based reference genome for A. *thaliana* on chromosome 1 (top) with the linear reference genome represented by blue nodes (bases). The figure shows a peak found for the ERF115 transcription factor that matches a DNA-binding motif. The peak detected by Graph Peak Caller follows the linear reference genome (blue nodes) except for the first SNP shown (green node), where the peak follows the green G instead of the blue A, making this a significant match against the DNA-binding motif (shown as a sequence logo at the bottom). The peak detected by MACS2 does not have a significant motif match.

For all transcription factors, except ERF115, the peaks uniquely found by Graph Peak Caller (not overlapping peaks found by MACS2) are more enriched for DNA-binding motifs than the peaks uniquely found by MACS2. In aggregate, the ratios of motif-matches of the uniquely found peaks are 7.04% for Graph Peak Caller and 6.03% for MACS2, yielding a *z*-value for the difference of 3.49 and a *p*-value of 0.024% using a one-sided *z*-test for difference in population proportions [17]. MACS2 in aggregate predicts 2443 more unique peaks than Graph Peak Caller, but only finds 11 (0.45%) more peaks that are motif-enriched, which is less than one would expect by chance, since the false discovery rate of motif-matches is controlled at 5%. Fig 3 shows one of the cases on chromosome 1 where Graph Peak Caller detects a peak that is not detected by MACS2, due to that the input reads are aligned against an indel that is not part of the linear reference genome.

**Fig 3.**
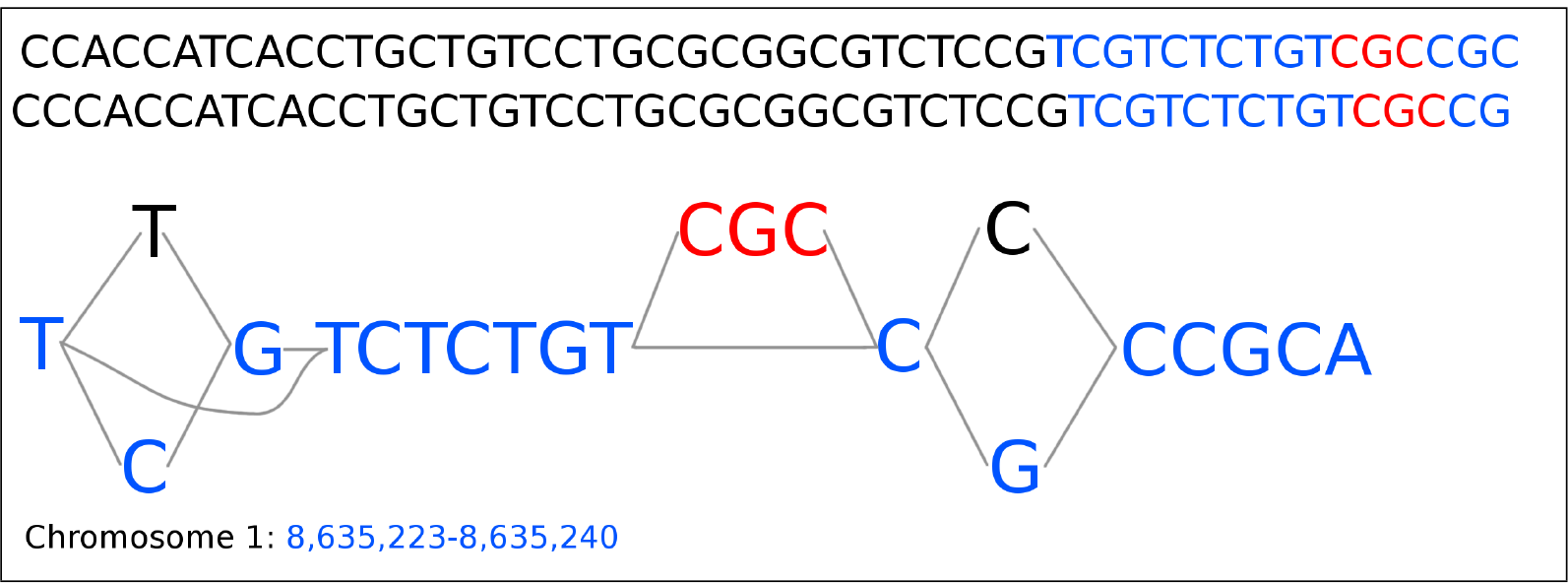
Example of part of a peak detected by Graph Peak Caller and not by MACS2. Showing part of the graph-based reference genome for *A. thaliana* on chromosome 1, containing two SNPs (black/blue) and an indel (red) that are not part of the linear reference genome (blue). From the raw ChIP-seq data for the ERF115 transcription factor (NCBI SRA SRR931836) two reads (shown at the top) align perfectly to the graph-based reference genome and Graph Peak Caller is able to detect a peak in this area. Mapping to the linear reference genome does not give sufficiently high mapping score, and so the peak is missed by MACS2.xs

The peaks found uniquely by Graph Peak Caller have more than twice the number of basepairs not part of the linear reference genome, compared to the peaks found by Graph Peak Caller that also have been found by MACS2 (Table S2). Fig 4 shows the proportion of peaks enriched for DNA-binding motifs for the peaks that are uniquely found by each peak caller (for a similar plot including all peaks, see Fig S1). As seen in the figure, Graph Peak Caller has a better correspondence between high peak scores and motif enrichment for all transcription factors, except ERF115.

**Fig 4.**
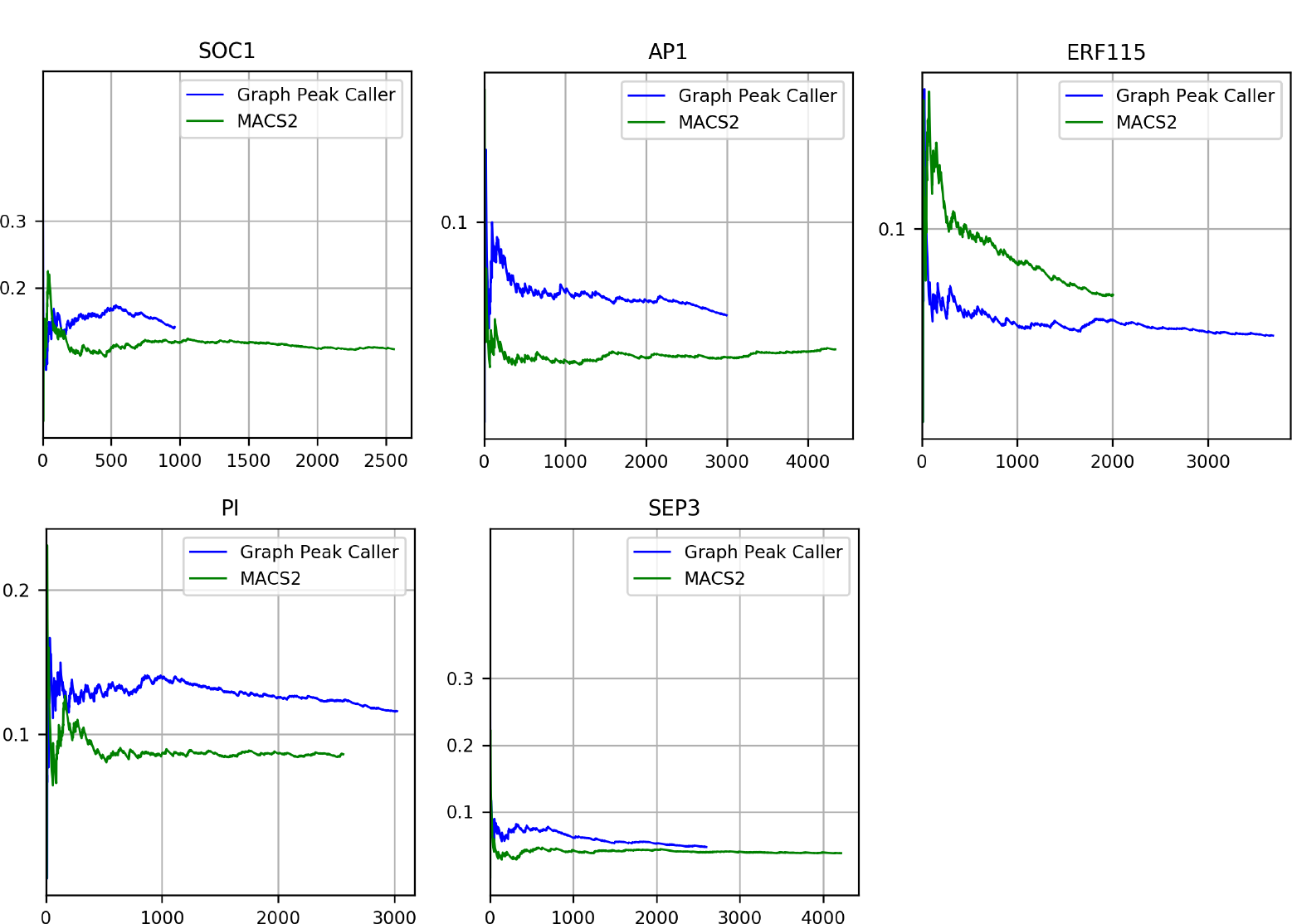
Dna-binding motif enrichment plots. The proportion of peaks enriched for a DNA-binding motif (Y-axis) when iteratively including more peaks from the set of uniquely found peaks for each peak caller, sorted descending on score (X-axis).

To validate that our peak caller works on other species, we repeated the experiment on similar datasets for *Drosophila melanogaster* and human (see Fig S2 and Table S1). In both cases Graph Peak Caller got a higher ratio of motif-matches on uniquely found peaks, but not statistically significant (*p*_human_ = 7%, *p*_D.melanogaster_ = 17%).

## Discussion

We have presented a first peak caller for ChIP-seq data mapped to a graph-based reference genome. We have tested our method by calling peaks on a graph-based reference genome for *A. thaliana* and comparing the detected peaks to those found by MACS2 using a linear reference genome of the same species. In the instances where the peak callers find peaks that overlap, Graph Peak Caller is able to find peaks that are more enriched for motifs. This is likely because Graph Peak Caller can trace variants within the peaks that are not part of the linear reference genome. Even if such peaks are also found by MACS2, the specific sequence comprising the DNA-binding motif may not be part of the linear reference.

Furthermore, Graph Peak Caller has a significantly higher proportion of motif-enriched peaks among its uniquely found peaks, showing that Graph Peak Caller is finding peaks enriched for motifs that MACS2 is not finding. These peaks covers more variations from the linear reference genome compared to the peaks found by both peak callers, and thus seem to be in areas where the advantages of graph-based reference genomes are more pronounced.

We chose to develop Graph Peak Caller by tightly following the principles of MACS2, so we easily could validate our graph-based approach and accurately measure the benefits of doing peak calling on a graph-based reference genome rather than on a linear reference genome. Having this as a working first approach to graph-based peak calling, it will now be natural and interesting to extend our work by drawing ideas from other peak callers or develop new peak calling principles to further improve graph-based peak calling. Also, Graph Peak Caller currently assumes no known information about the specific paths of the diploid genome of the individual that ChIP-seq data has been collected from. It would be interesting to develop a ChIP-seq pipeline where the path(s) through the reference graph are known (or estimated based on the ChIP-seq data), and compare that approach to Graph Peak Caller.

There are a few challenges with a graph-based ChIP-seq approach. Mapping to graphs is still in its infancy, and has not yet reached its full potential. Also, both mapping to graph-based reference genomes as well as many of the operations required for doing peak calling, such as expanding input reads, are, in our experience, still a lot slower than existing solutions on linear reference genomes.

We believe that our peak caller represents an important step towards creating a more comprehensive toolset for functional genomics on graph-based reference genomes, extending the possible applications of graph-based reference genomes and bringing the genomics community an important step closer to widespread adoption of these reference structures.

## Methods

### Peak calling

Our approach to graph-based peak calling is implemented in an open source Python 3 package, *Graph Peak Caller*. Graph Peak Caller was developed by extending the methodologies and concepts from MACS2 to directed acyclic graphs (DAGs). The MACS2 algorithm can be divided into five steps: estimating the fragment length, creating a fragment pileup by extending input reads to match the estimated fragment length, calculating a background track based on local and global average number of reads, calculation of p/q scores based on the fragment pileup and background track, and finding peaks based on thresholded scores. We have adopted each of these steps to work on DAGs. Fig 1 illustrates the method on a graph-based reference genome, and the following describes the details of each step.

Graph Peak Caller uses the linear estimation algorithm from MACS2 to estimate the fragment length *f* by using the linear path through the graph with the highest number of aligned reads as reference. Graph Peak Caller generates *the fragment pileup* by extending each read to the estimated fragment length *f*, and counting the number of extended reads that cover each base pair in the graph. For a single read with length *r*, the extension is done by including all possible paths of length *f-r* in the graph that start at the read’s end position. The background track is an estimate of the expected number of reads mapping to each position in the reference. This is, for a given position in the reference, estimated by measuring the amount of reads mapping in the “neighbourhood” of that position. The reads can either be the input reads or a set of control reads. On a linear reference genome, the background track is simply estimated by taking the average pileup count in a local window around each base pair. This is less trivial to do on a graph-based reference genome, since the concept of neighborhood is not as well defined. We solve this problem by projecting the graph onto a single linear path where parallel paths are projected to the same position on the linear path. This allows us to perform background track estimation much the same way as MACS2 does, using a linear reference, and then projecting the resulting track back to the graph again. If control reads are used to generate the background track, the background track is scaled with the ratio of control reads to input reads.

The fragment pileup and background track are then treated as counts and rates in Poisson distributions, and p-values are computed for each position for the observed *count*, given the corresponding *rate*. Since one test is performed for each position in the graph, we compute q-values (adjusted p-values) to control the false discovery rate. The q-values are thresholded at a user specified threshold, yielding a binary track of potential binding regions.

Graph Peak Caller then removes small gaps (similarly to MACS2) between these potential binding regions. On a graph, this is done by joining regions that are connected by a path shorter than the read length. If a gap consists of several paths, all paths of length shorter than the read length is included in the joined region. Then, the resulting regions are grouped into connected subgraphs, representing areas of potential binding events. The final peaks are selected by finding the path through each subgraph that has the highest number of input reads mapped to it. Similarly to MACS2, peaks that are shorter than the estimated fragment length are removed.

For each subgraph, Graph Peak Caller can also report an “alternative” peak in addition to the main peak. This is done by using Fimo [10] to estimate the exact location within the peak subgraph that matches the binding motif, and looking for an alternative path through this area which is covered by at least one input read. Such alternative peaks can be used to infer differential binding.

### Validation and testing

To test our peak caller, we used *vg* [6] to create a whole genome *Arabidopsis thaliana* reference graph by using variants from the 1001 Genomes Project. We selected all transcription factors listed in the transcription factor database of *Expresso* [11] that also had a motif in the *Jaspar* database of transcription factor binding profiles [12], resulting in a set of 5 transcription factors: ERF115, SEP3, AP1, SOC1, and PI. (Two transcription factors, SVP and ATAF1, were omitted due to invalid fastq files. AP2 and AP3 were omitted based on their close relatedness to AP1. Also, PIF3 was omitted since neither the detected binding events by Graph Peak Caller nor MACS2 had any association with the motif we found in the Jaspar database). Raw ChiP-seq reads were downloaded from the NCBI Sequence Read Archive (SRA) (SRA accession numbers in Appendix S1) and trimmed using *Trim Galore!* v0.4.4 [13] (default parameters). Reads were mapped both to our graph-based reference genome using *vg* and to the *Tair10* [15] reference genome using *BWA* v0.7.12 (`bwa aln followed by bwa samsse`, default parameters). In the linear case we filtered out low-quality alignments using *SAMtools* v0.1.19 [14] with the command samtools `view −F 1804 −q 37`, and for the graph alignments we used `vg filter −q 37`. *MACS2* v2.1.0 was used to call peaks on the linear reference genome, using default parameters. We created DNA-binding motif enrichment plots (Fig 4) for each set of detected peaks (URLs to the motif models that were used are in Appendix S1). We have created a Docker repository with the *A. thaliana* graph-based reference genome, Graph Peak Caller, *vg* and all other software and scripts used to generate the results in this article. A simple guide on how to re-run the experiments can be found in the Github repository for Graph Peak Caller.

## Conclusion

We have developed Graph Peak Caller, a tool for performing peak calling from ChIP-seq reads mapped to a graph-based reference genome. Graph Peak Caller is based on the same principles as MACS2. We have validated our approach by using both Graph Peak Caller and MACS2 to call peaks using ChIP-seq datasets on *A. thaliana*, showing that the peaks found by Graph Peak Caller in general are more enriched for DNA-binding motifs than those found by MACS2 on a linear reference genome. Graph Peak Caller is also able to provide candidates for differentially expressed peaks, and it provides a first method for doing peak calling on graph-based reference genomes.

## Supporting information

S1 Table Overview of peaks detected on human and *D. melanogaster*.

S2 Table Average number of base pairs not part of linear reference genome.

**S1 Fig. DNA-binding motif enrichment plots (as Fig 4) for all peaks detected on *A. thaliana***. Contrary to Fig 4, these plots include all peaks found by both peak callers.

**S2 Fig. DNA-binding motif enrichment plots (as Fig 4) for all peaks detected on *D. melanogaster* and human.** Left plots are proportion of peaks matching motif when all peaks are included. Right plots are proportion of peaks matching motif when only unique peaks found by each peak caller are included.

S1 Appendix. URLs to motifs and accession numbers to data used in experiments.

## Acknowledgements

The sequence data used for creating the *Arabidopsis thaliana* reference graph were produced by the Weigel laboratory at the Max Planck Institute for Developmental Biology.

## References

1. Zhang Y, Liu T., Meyer CA, Eeckhoute J., Johnson DS, Bernstein BE., Nusbaum C, Myers RM., Brown M, Li W., Liu XS. Model-based Analysis of ChIP-Seq (MACS). Genome Biology. 2008 Nov;9(9):R137.

2. Kharchenko PV, Tolstorukov MY., Park PJ. Design and Analysis of ChIP-seq Experiments for DNA-binding Proteins. Nature Biotechnology. 2008 Dec;26(12):1351.

3. Wilbanks EG, Facciotti MT. Evaluation of Algorithm Performance in ChIP-seq Peak Detection. PloS One. 2010 Jul 8;5(7):e11471.

4. Paten B, Novak AM., Eizenga. JM, Garrison E. Genome Graphs and the Evolution of Genome Inference. Genome Research. 2017 May 1;27(5):665–76.

5. Novak AM, Hickey G., Garrison E, Blum S., Connelly A, Dilthey A., Eizenga J, Elmohamed MS., Guthrie S, Kahles A., Keenan S. Genome Graphs. bioRxiv. 2017 Jan 1:101378.

6. Garrison E, Sirén J, Novak AM, Hickey G, Eizenga JM, Dawson ET, Jones W, Lin MF, Paten B, Durbin R. Sequence Variation aware Genome References and Read Mapping with the Variation Graph Toolkit. bioRxiv. 2017 Jan 1:234856.

7. Li H. Aligning Sequence Reads, clone Sequences and Assembly Contigs with BWA-MEM. arXiv preprint arXiv:1303.3997. 2013 Mar 16.

8. Langmead B, Salzberg SL. Fast Gapped-read Alignment with Bowtie 2. Nature Methods. 2012 Apr;9(4):357.

9. Rand KD, Grytten I., Nederbragt AJ, Storvik GO., Glad IK, Sandve GK. Coordinates and Intervals in Graph-based Reference Genomes. BMC Bioinformatics. 2017 Dec;18(1):263.

10. Grant CE, Bailey TL., Noble WS. FIMO: Scanning for Occurrences of a given Motif. Bioinformatics. 2011 Feb 16;27(7):1017–8.

11. Aghamirzaie D, Velmurugan KR., Wu S, Altarawy D., Heath LS, Grene R. Expresso: A Database and Web Server for exploring the Interaction of Transcription Factors and their Target Genes in Arabidopsis thaliana using ChIP-Seq peak Data. F1000Research. 2017;6.

12. Sandelin A, Alkema W., Engström P, Wasserman WW., Lenhard B. JASPAR: an open-access database for eukaryotic transcription factor binding profiles. Nucleic acids research. 2004 Jan 1;32:D91–4.

13. Krueger F. Trim galore. A Wrapper Tool around Cutadapt and FastQC to consistently apply Quality and Adapter trimming to FastQ files. 2015.

14. Li H, Handsaker B., Wysoker A, Fennell T., Ruan J, Homer N., Marth G, Abecasis G., Durbin R. The Sequence Alignment/Map Format and SAMtools. Bioinformatics. 2009 Aug 15;25(16):2078–9.

15. Arabidopsis Genome Initiative. Analysis of the Genome Sequence of the flowering Plant Arabidopsis thaliana. Nature. 2000 Dec;408(6814):796.

16. 1000 Genomes Project Consortium. A global Reference for Human genetic Variation. Nature. 2015 Oct;526(7571):68.

17. Devore JL, Berk KN. Modern Mathematical Statistics with Applications. 2nd ed. New York: Springer-Verlag; 2012. p. 519–521

