## Supplementary Materials for "Graph Peak Caller: calling ChIP-Seq Peaks on Graph-based Reference Genomes" (from another version of this preprint: version 1, 2018-03-23)

### S1 appendix

#### Reference graphs

All graphs used in the experiments can be downloaded from:

- *Arabidopsis thaliana: hyperbrowser.uio.no/graph-peak-caller/static/graph_peak_caller_data/arabidopsis_thaliana.tar.gz*
- *Drosophila melanogaster: hyperbrowser.uio.no/graph-peak-caller/static/graph_peak_caller_data/drosophila_melanogaster.tar.gz*
- Human: hyperbrowser.uio.no/graph-peak-caller/static/graph_peak_caller_data/human.tar.gz

#### ChiP-seq reads and motifs used

For all transcription factors on *Arabidopsis thaliana*, raw ChIP-seq reads were fetched from the NCBI Sequence Read Archive using the following accession numbers:

- ERF115: SRR931836
- SEP3: SRR1042995
- AP1: SRX387187
- SOC1: SRR1044950
- PI: SRR502859

For the validation on human and *Drosophila melanogaster,* raw ChIP-seq reads were collected from experiments on ENCODE having the following experiment IDs. In all cases, raw fastq reads from replicate 1 were used:

- Human CTCF: ENCSR000DUB
- Human SRF: ENCSR000BIV
- *D. melanogaster* JRA: ENCSR471GSA
- *D. melanogaster* SQZ: ENCSR923VWW
- *D. melanogaster* ANTP: ENCSR082RBU
- *D. melanogaster* JIM: ENCSR978WED

The motifs were collected from the Jaspar database [1] (for human and *A. thaliana*) or from OnTheFly [2]:

- *A. thaliana* ERF115*: https://hyperbrowser.uio.no/graph-peak-caller/static/graph_peak_caller_data/motifs/ARABIDOPSIS_ERF115.meme
- *A. thaliana* SEP3: http://jaspar.genereg.net/api/v1/matrix/MA0563.1.meme
- *A. thaliana* AP1: http://jaspar.genereg.net/api/v1/matrix/MA0940.1.meme
- *A. thaliana* SOC1: http://jaspar.genereg.net/api/v1/matrix/MA0554.1.meme
- *A. thaliana* PI: http://jaspar.genereg.net/api/v1/matrix/MA0559.1.meme
- Human CTCF: http://jaspar.genereg.net/api/v1/matrix/MA0139.1.meme

- Human SRF: http://jaspar.genereg.net/api/v1/matrix/MA0083.2.meme
- *D. melanogaster* JRA: https://hyperbrowser.uio.no/graph-peak-caller/static/graph_peak_caller_data/motifs/DM_JRA.meme
- *D. melanogaster* SQZ: https://hyperbrowser.uio.no/graph-peak-caller/static/graph_peak_caller_data/motifs/DM_SQZ.meme
- *D. melanogaster* ANTP: https://hyperbrowser.uio.no/graph-peak-caller/static/graph_peak_caller_data/motifs/DM_ANTP.meme
- D. melanogaster JIM: https://hyperbrowser.uio.no/graph-peak-caller/static/graph_peak_caller_data/motifs/DM_JIM.meme

* Since we were not able to find a motif for A. thaliana ERF115 in the Jaspar database, we used the following motif from the Plant Cistrome Database [3]: http://neomorph.salk.edu/dap_web/pages/browse_table_aj.php?AGI=ERF115

#### References

[1] Khan, A. et al. JASPAR 2018: update of the open-access database of transcription factor binding profiles and its web framework. Nucleic Acids Res. 2018; 46:D260–D266, doi:10.1093/nar/gkx1126

[2] Shazman, Shula, Hunjoong Lee, Yakov Socol, Richard S. Mann, and Barry Honig. "OnTheFly: a database of Drosophila melanogaster transcription factors and their binding sites." Nucleic acids research 42, no. D1 (2013): D167-D171.

[3] Ronan C. O'Malley, Shao-shan Carol Huang, Liang Song, Mathew G. Lewsey, Anna Bartlett, Joseph R. Nery, Mary Galli, Andrea Gallavotti, and Joseph R. Ecker (2016). Cistrome and Epicistrome Features Shape the Regulatory DNA Landscape, Cell. 2016 May 19;165(5):1280-92. doi: 10.1016/j.cell.2016.04.038.
