## Supplementary figures and images for "Graph Peak Caller: calling ChIP-Seq Peaks on Graph-based Reference Genomes" (from another version of this preprint: version 1, 2018-03-23)

### Supplementary Materials

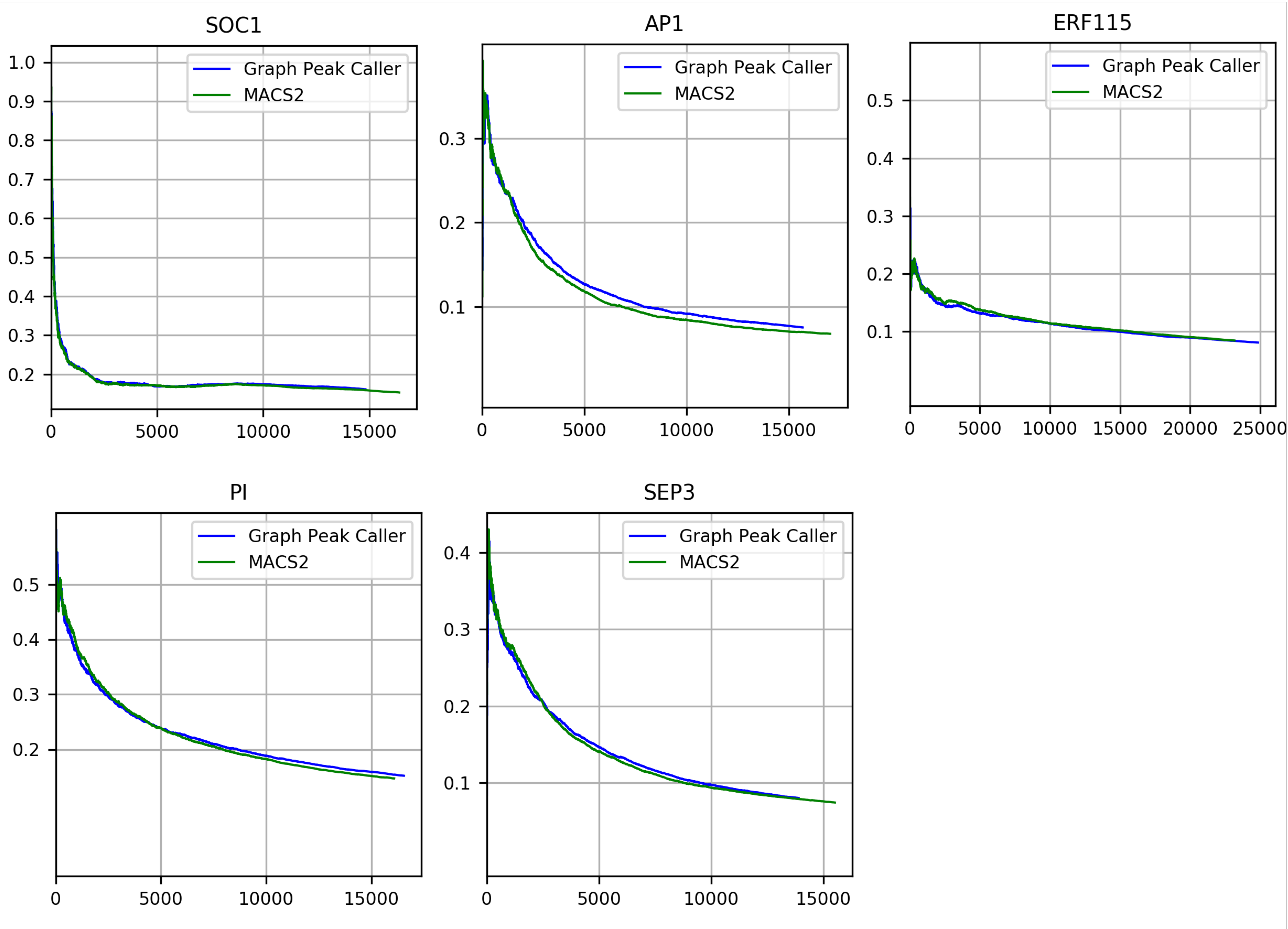

### Supplementary Materials

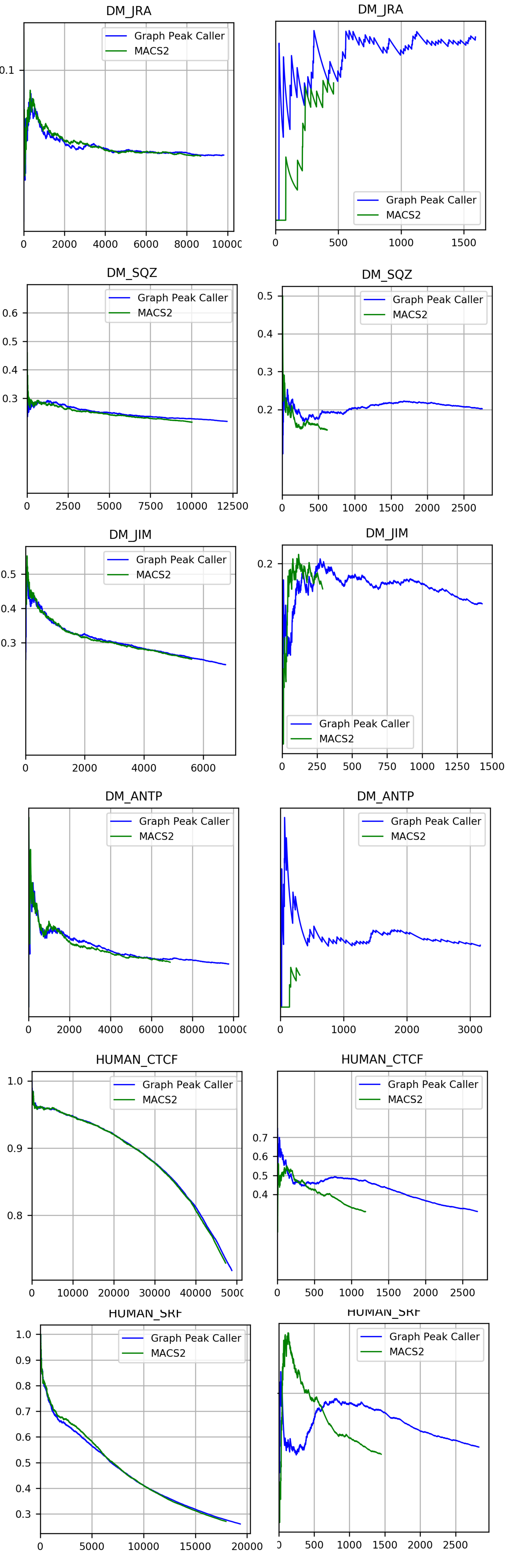
